# Topological gene-expression networks recapitulate brain anatomy and function

**DOI:** 10.1101/476382

**Authors:** Alice Patania, Pierluigi Selvaggi, Mattia Veronese, Ottavia Dipasquale, Paul Expert, Giovanni Petri

## Abstract

Understanding how gene expression translates to and affects human behaviour is one of the ultimate aims of neuroscience. In this paper, we present a pipeline based on Mapper, a topological simplification tool, to produce and analyze genes co-expression data. We first validate the method by reproducing key results from the literature on the Allen Human Brain Atlas, and the correlations between resting-state fMRI and gene co-expression maps. We then analyze a dopamine-related gene-set and find that co-expression networks produced by Mapper returned a structure that matches the well-known anatomy of the dopaminergic pathway. Our results suggest that topological network descriptions can be a powerful tool to explore the relationships between genetic pathways and their association with brain function and its perturbation due to illness and/or pharmacological challenge.

## Introduction

The human brain is a highly complex organ where its function emerges from the integration of cellular, anatomical and functional circuits [3]. This complexity is thought to be crucial to the plasticity needed to adapt to environmental changes. The architecture of the human brain is ultimately shaped by the human genome through the regulation of gene expression. In fact, the human brain consists of a set of differentiated regions each having a specific distribution of cell types and a microscopic and macroscopic anatomical organization that are the results of the differential expression of unique gene patterns during development that are kept stable when maturity is reached [16]. Traditionally, genetic studies have investigated the association of genetic variants with a variety of brain disorders in large population studies [28, 35]. However, these studies are not particularly informative on the impact of these gene variants on brain structure and function. Imaging-genetic studies provided additional insights by exploring the effect of genetic variants and expression of gene sets on normal and pathological brains. They are however not without limitations [4]; a crucial one is that they look at the association between gene network and brain phenotype in a limited number brain regions, e.g. the prefrontal cortex. However, the availability of new data-sets such as the Allen Human Brain Atlas data set (AHBA) [13], with vastly enlarged brain coverage and resolution, offered a unique opportunity to explore the architecture of the brain transcriptomics in the entire brain. Critically, such a comprehensive data set allows the investigation of the architecture of differential gene expression between brain regions. In fact, works that explored the complexity of the human brain transcriptome, taking advantage of the AHBA data set, are starting to uncover new knowledge about normal and pathological brain function. For example, [14] revealed large transcriptional differences between brain regions. In addition, the same authors analyzed the expression of genes with patterns of differential expression across regions that are highly consistent between donors, a measure they called differential stability showing that: i) these patterns reflects the physical topography and the developmental trajectories of brain regions, ii) these genes are strongly associated with different brain disorders, and iii) the expression of these genes in the cortex is correlated with resting state functional connectivity. Also, other authors found that resting state functional connectivity networks are associated with between-brain-tissues correlated expression of a set of 136 genes [27]. Both studies consolidated early evidence that mesoscale differences and between brain regions similarities in terms of structure and function find their backbone in the architecture of the human transcriptome. Network approaches have been extensively used to investigate the complexity of the brain by using a variety of brain imaging data including Magnetic Resonance Imaging (MRI), Electroencephalography (EEG) and Magnetoencephalography (MEG). These studies revealed that some topological properties of the brain are highly conserved across different scales and type of measures [6]. In particular, the brain appears as a topological structure characterized by short path length and high clustering, organized in hubs and shaped in modular communities [5]. Assuming that the brain is a self similar system [12, 34], we expect that its transcriptomics could be described with topological analysis. Topological characteristics of the brain transcriptome have been already investigated by some authors [18, 30]. However, one key limitation in the analysis of the topology of the brain transcriptomic is the high-dimensionality of the data. The AHBA features the expression levels of *>*20,000 genes from six post-mortem brains, profiled by *∼* 60,000 microarray probes in different brain regions that are spatially-resolved at different level of anatomical coarseness [13]. As a result, the analysis of the AHBA data set poses a significant problem of high-dimensionality that can be tackled by using different strategies in variable selection and feature extraction [15]. Here we present a new approach based on the Mapper algorithm [33] to reduce the dimensionality of microarray mRNA expression data from the AHBA while preserving topological information and characteristics of the human brain transcriptome. The Mapper algorithm is a tool that has been developed to identify topological characteristics of data sets based on the distance between data points after applying a pre-definite filter function [33]. The algorithm constructs a series of networks describing the data-set at different levels of coarseness and, by using the composition of each node to map the samples to their anatomical location, we can represent the similarity of the genetic expression of samples obtained from different ROIs as network connectivity patterns. Mapper has been already successfully used to analyse high dimensional behavioural, clinical, biological and neuroimaging data sets [19, 20, 24, 31]. In this work, we present three different applications of the Mapper in the microarray AHBA data set: i) replication of the gene co-expression analysis originally presented by the Allen Institute for Brain Science [13]; ii) topological co-expression analysis of the gene list identified by [27]; iii) topological co-expression analysis of the genes in the dopamine pathway. The aim of these three case-studies is first to validate the pipeline by replicating previous findings and then to test its ability to extract useful information from the dopamine system that is known to be crucial for many brain disorders.

## Results

In this section, we first briefly introduce the Mapper algorithm and the agreement matrix analysis we used (a more in depth description can be found in the Methods section) before introducing the data sets we analyzed. Finally, we present and discuss in details the results we obtained on the three data set we considered.

### Gene Mapper networks and agreement matrices

The Mapper algorithm was first introduced in [33] as a technique to extract low-dimensional skeletons for the classification of 3D shapes. In recent years, however, its use as a data analytic tool has vastly extended. In its simplest form, the algorithm takes as input a set of data points equipped with a similarity metric and returns a network which encodes a low-dimensional backbone of the data set that can be interpreted as a network. In the present study, we use a correlation-based distance to group together the gene-expression vectors of brain tissue samples from the left hemisphere of six donors [1]. We use the first two principal components, which explain between 21.6% and 30%, depending on the gene list considered, of the variance in the gene-expression to guide the algorithm in the local clustering, and we optimize the parameters in order to obtain a good ratio between noise, i.e. un-connected voxels, and signal, i.e. connected components, with we which we obtain a series of networks describing the data-set at different levels of coarseness. Details about the construction and robustness of the methods are given in the Supporting Information (S.I.). Each node in the network produced by Mapper represents a cluster composed by samples with very similar gene-expression profiles and similar loading on the first and second principal components. Therefore, we can represent the similarity of the genetic expression of samples obtained from different ROIs by mapping the samples constituting each node to map the samples to their anatomical location. The connectivity patterns between nodes in the network further highlight this similarity. In particular, we can study the subgraphs determined by nodes that contain samples from the same ROIs. Scattered subgraphs indicate a loose similarity within the specific ROI and a higher similarity with other ROIs, see Fig.1 for an illustration. Given a set of parameters *{σ}* for the Mapper, see Methods for details, this information can be further summarized using a co-occurrence matrix *A*[*σ*], where each element *A_ij_*[*σ*] counts the numbers of times a sample from region *i* and a sample from region *j* are mapped to the same node. We obtain a co-occurrences matrix for each optimal set of parameters and further compress the information obtained from the networks corresponding to different parameter choices in an agreement matrix *A* = *〈A*[*σ*]*〉_σ_*, computed by averaging across all matrices and retaining the non-zero elements that have at least one connection in all the networks.

**Figure 1:**
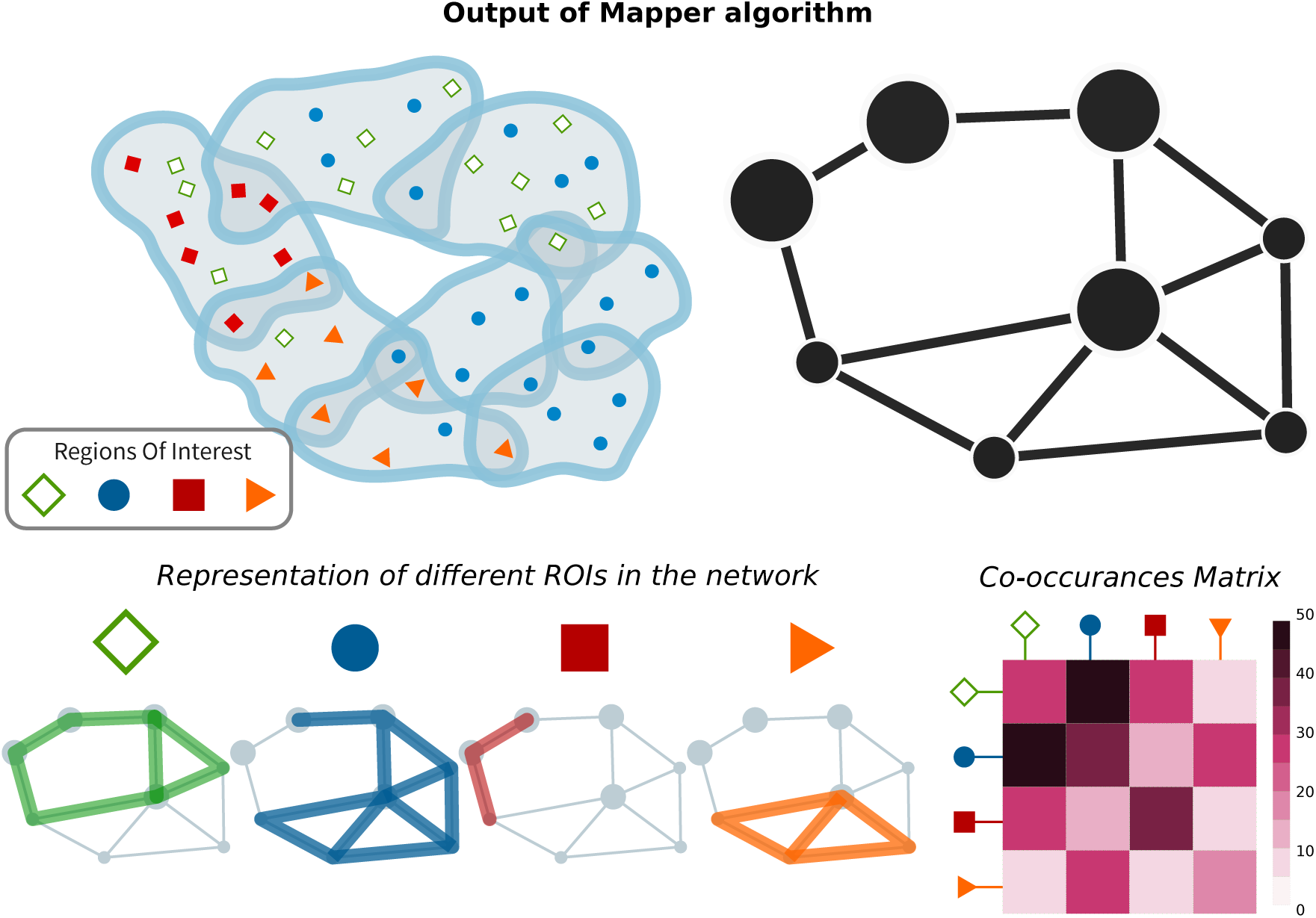
Representations and Analytical Tools for the Mapper output. After slicing the data space in overlapping slices, partial clustering is applied to each slice (a). Since some points belong to adjacent slices, they will belong to clusters in both slices (a). These overlapping clusters effectively produce a cover of the dataspace and can be summarised as nodes (containing the data points) linked to other nodes, whenever two clusters share data points (b). Data points correspond to samples of brain regions with specific anatomical and spatial characterizations. It is therefore possible to investigate how distributed or –conversely– localized the samples of a certain Region of Interest (ROI) are (c). This information is succinctly described by the co-occurrences matrix that counts how often nodes belonging to different ROIs belong to the same Mapper node (d).

### Data sets

We applied the construction described above to three different sets of genes: the whole human genome (*∼* 29000 genes), a list of 136 genes that support synchronous activity in brain networks as shown by [27], and a list of 56 genes related to the dopamine system. The dopamine list was created by interrogating the Gene Ontology database (http://geneontology.org) and the Panther gene classification system [22] from which the “Dopamine receptor mediated signaling pathway (P05912)” list was selected. The gene expression data for all three gene-lists we considered come from the microarray data of the Allen Human Brain Atlas (http://human.brain-map.org). The original log2 data were transformed into z-scores following the methodology present in [29]. We use the first data set to validate the proposed methods within a fully gene-focused setup. The analysis of the second dataset instead highlights the capacity of this method to meaningfully link the genetics and activation level of specific ROIs. Finally the dopamine system analysis provides new insights in brain organisation. We used both the agreement matrices and standard network properties, e.g. shortest path distances, to compare our results with previous analysis done on the same data set. Functional connectivity matrix was computed using high-resolution resting-state fMRI data from 20 subjects (5F/5M aged 26–30; 5F/5M aged 31–35) were randomly selected as part of the Human Connectome Project (h2ps://db.humanconnectome.org/). Please find full details about data acquisition and pre-processing in S.I.)

### Validation against the Allen Brain Atlas

Fig. 2 shows a few examples of Mapper network (parameters q=5, overlap = 20, 25, 35, see Methods) obtained for the Allen Brain Atlas (AHBA) data set at different levels of coarseness. We can see that in all cases few major connected components in the network: a small one, containing only elements of the cerebellum, a giant component containing mostly cortical samples, and smaller subcortical components. As the overlap increase, we can see how the larger components starts developing connections towards the components containing samples from subcortical ROIs, while the component containg samples from the cerebellar cortex is still very much isolated in all cases.

**Figure 2:**
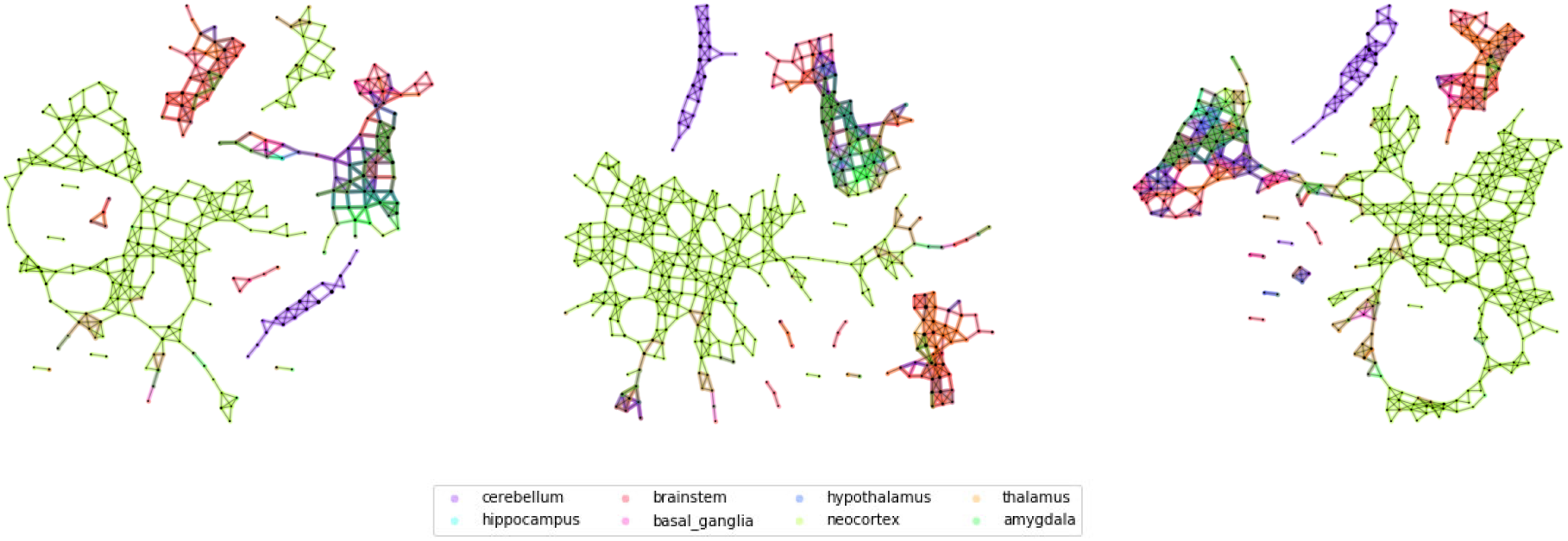
Mapper networks of the Allen Brain Atlas data set. From left to right, we show some Mapper networks obtained for the same size but different overlaps between windows (overlap 25%, 30%, 35%). The networks display very similar qualitative properties, e.g. the separation of the cerebellar areas, which we further characterize using the agreement matrices.

While some of these patterns and their stability can be guessed by direct observation, the distinction is much clearer when we consider the structure of the agreement matrix (Fig. 3a) (for parameters (window size, overlap)=*{*(5, 25),(5, 30),(5, 35),(6, 20)*}*, See S.I. for more information on the parameter choice) where we can clearly distinguish 4 blocks representing: the cerebral cortex, the hippocampus, the cerebellum, and brainstem nuclei. We can see some regions of the cerebral cortex that have higher connectivity with the two blocks from the subcortex, indicating the connection in the network of the central community to the peripheral structures. We are now in a position to compare the findings of Mapper to the original paper [13]. We do this by comparing agreement matrices for the Mapper network with the co-expression matrix of the original paper, which is summarised in Fig.3. To guide the eye, we ordered the regions in Fig. 3 in the same order as in [13] (Fig. 3b) and it is easy to see that the Mapper agreement matrix reproduces the general structure of the differential gene-expression matrix. In order to quantify this effect, we delve a little deeper in the relationship between Mapper links and the standard differential gene-expression techniques. In fact, we show that Mapper extracts the connections with lower differential of gene-expression. To do this, we compare the differential gene-expression between the ROIs that are connected in the Mapper and the ones that are not. In particular, we expect that present links should be characterized by smaller differential expressions, since the presence of the link suggests a higher similarity between the samples contained in the nodes, while the absence of a link supports the opposite. To quantify this difference we use the Kolmogorov–Smirnov statistic which measures a distance between the empirical distribution functions of two samples (Fig. 3e). We find that links present in the Mapper generated networks are up to 27 % more likely to include low differential co-expression between the ROIs than the complete full matrix, and up to 45% more likely than the ones ignored by the algorithm construction (3c). When confronting the two differential co-expression distributions for present and absent links (3d) we can clearly see that Mapper tends to contain more links with a differential gene-expression of at most *e*^4.85^ than the ones it excludes.

**Figure 3:**
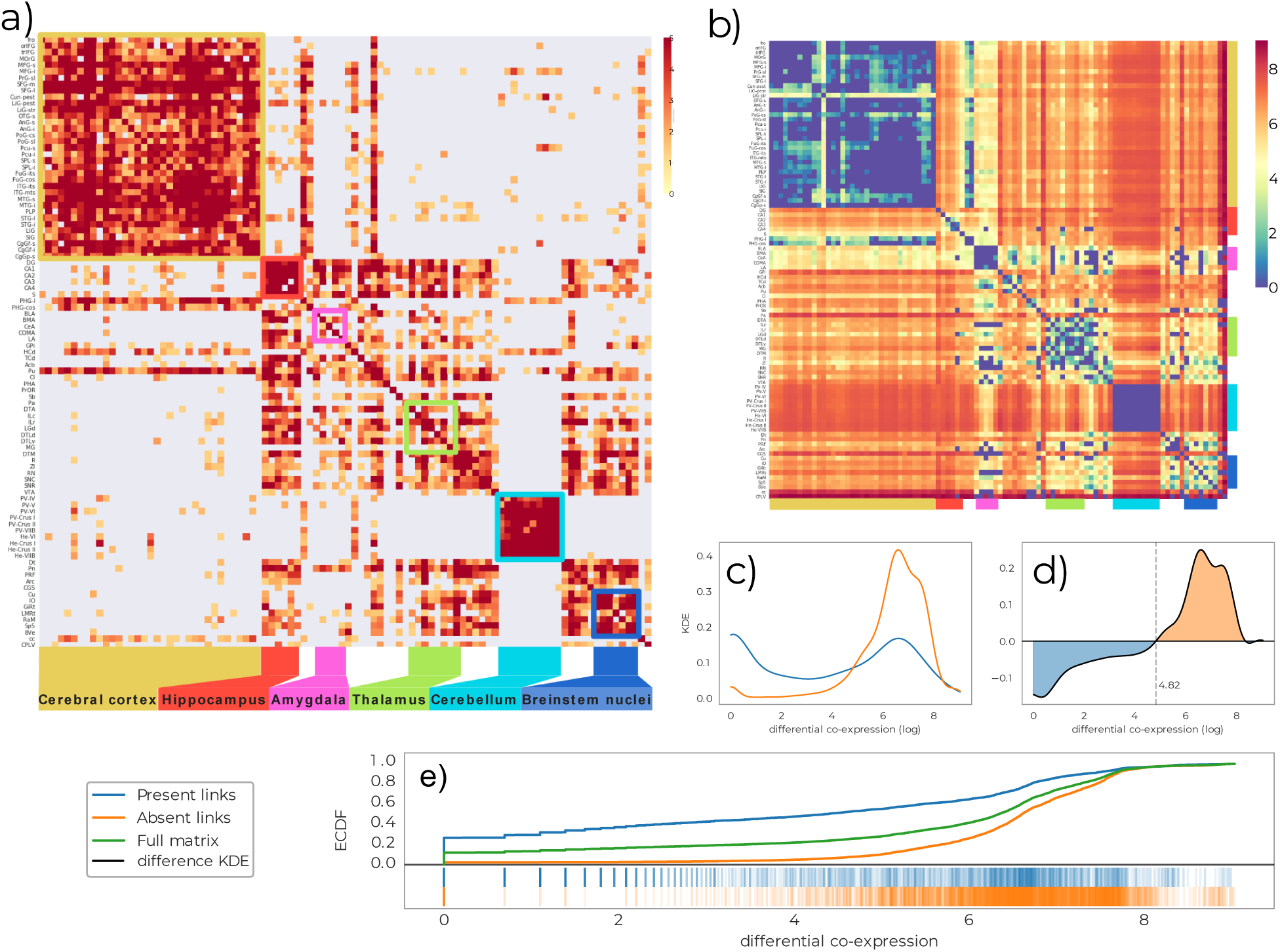
Comparison of the output network with differential analysis from the Allen Brain Institute. a) Agreement matrix between Mapper co-occurence matrices for parameters (window size, over-lap)=*{*(5, 25),(5, 30),(5, 35),(6, 20)*}* (See S.I. for more information on the parameter choice). b) Differential gene-expression matrix, reproduced from [14]. c) distributions of (log) differential gene-expression for links that are and are not present in the Mapper networks. d) difference between the distributions in c). e) cumulative distributions.

In addition to the large-scale results reported above, it is possible to focus on the comparison of the brain systems analysed in [14]. In Fig. 4a, we reproduce the agreement matrix coloured according to four regimes. These are based on the combinations of sparse/dense connectivity in the Mapper and of low/high differential gene expression in the nodes. A link is qualified as dense if there are on average more than 2 co-occurances between the nodes, and the low/high differential expression is defined by the trade-off detected in Fig.3d. It is clear that the two most represented cases are the expected ones: few different genes expression and dense Mapper connectivity, in light blue, versus inconsistent genes expression profiles and sparse network connectivity, in dark red. In the S.I. we give more details about how the classification in four regimes is done.

**Figure 4:**
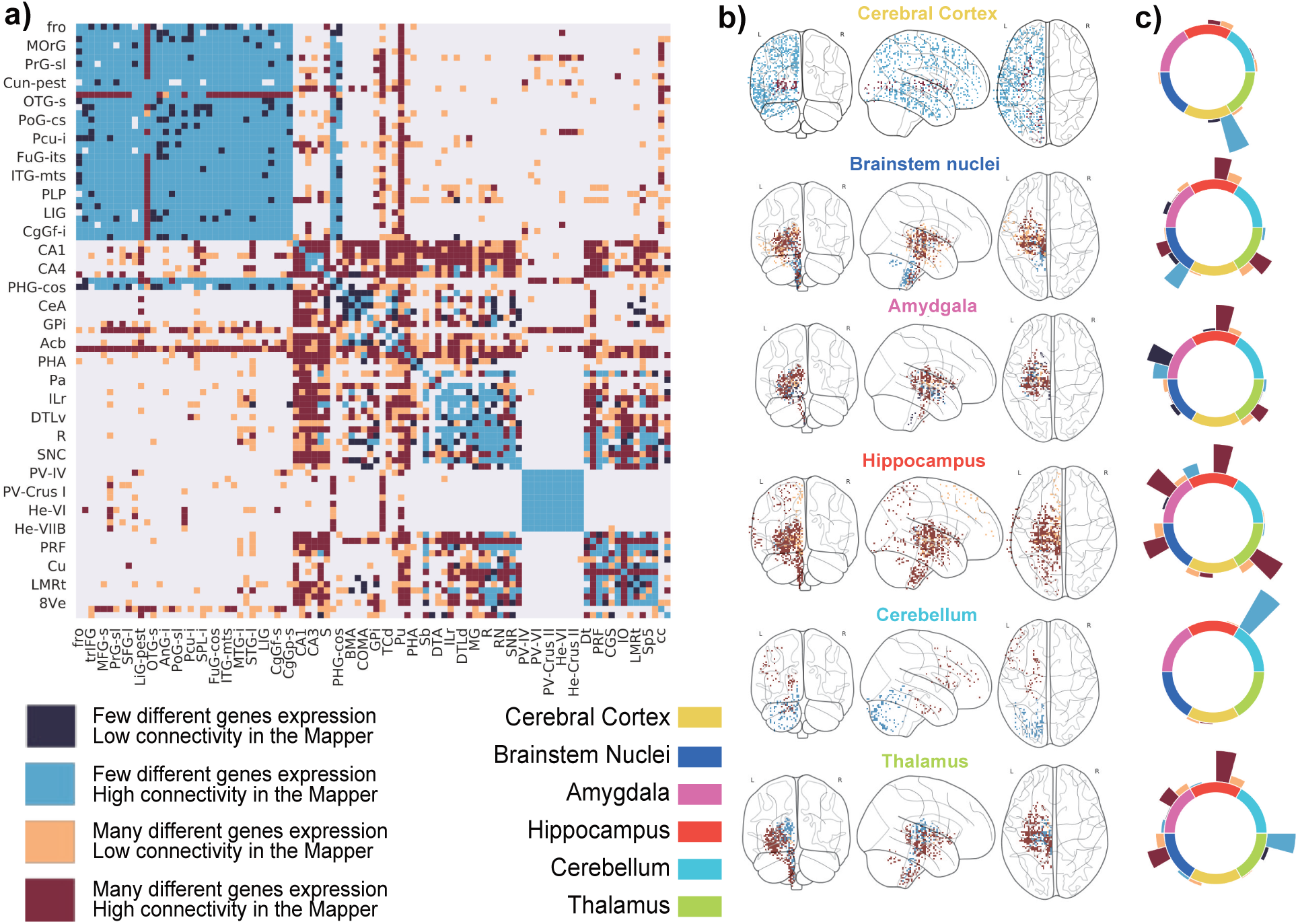
Comparison between the connectivity and the differential gene-expression for links present in the Mapper. a) The agreement matrix coloured according to four regimes based on the combinations of sparse/dense connectivity in the Mapper (defined as more or less than 2 co-occurances between the regions) and of low/high differential gene expression (defined using the trade-off detected in Fig.3d). b) and c) Visualizing the predominant regimes of the links connecting each region of interest with the rest of the hemisphere. In b) only samples belonging to areas connected with the ROI are shown colored according to their relative regime. In c) the we visualize the total number of links between the ROIs belonging to each regime.

Beyond this broad classification into 4 regimes, there are more subtle connectivity patterns: for example the cerebral cortex and the cerebellum are clearly more homogeneous than the hippocampus or the amygdala (Fig. 4b). To extract all the information from the Mapper results and interpret them correctly, we need to go further. We can quantify for each ROIs how many links of a certain type link to other ROIs. We summarise this construction in the circular barplots of Fig. 4c. In general, we can see how for low differential gene expression the Mapper tends to always be densely connected, while the viceversa is not always truw with clear examples in the hippocampus and amygdala.

Using this information we can compare the Mapper results with those of the AHBA for each system/ROI:

- **Cerebral Cortex:** The cerebral cortex appears to be very connected in the Mapper network, consistent with the idea that the basic architecture across the entire cortex is similar or “canonical”. This accordance of low differential genes expressed and high connectivity is indicated by the light blue shading comprising cortical gyri ordered from the frontal pole (fro) to cingulate gyrus (CgGP-s). The sole exception is the visual cortex LiG-str, which had a uniquely diverse gene-expression that the network was not able to discern.
- **Cerebellum:** The same connectivity pattern can be seen for the cerebellum. [14] noticed how the internal homogeneity across subdivisions of the cerebellum, with samples from different cerebellar lobes listed from “PV-IV” through “He-VIIB” show no internally differentially expressing genes. This peculiarity is clearly picked up by the Mapper outputs, as the cerebellum always creates a connected component of its own in the network.
- **Hippocampus:** In [14], the hippocampus showed a distinct pattern of gene expression across its highly distinct and stereotyped anatomical divisions. This high number of differential gene-expression is not reflected in the mapper generated network by a disconnection. Instead, the samples from the various hippocampal structure are highly connected within the subcortex and loosely connected with parts of the cortex (LiG-pest, SPL-s, SPL-i, STG-l). This pattern is different from its close relative the cortex, and even more distinct from that of evolutionary older brain regions.
- **Amygdala:** In their work, [14] found that the amygdala are very similar to one another while very different from other brain regions, in the mapper these regions are instead loosely connected within the subcortex and within itself.
- **Thalamus, Brainstem Nuclei:** The thalamus and brainstem nuclei show a great deal of complexity in the differential gene-expression matrix. This is confirmed in the mapper agreement matrix. We can see how these two structures are interconnected in the network but with high incidence of low connectivity links suggesting a less clear organization than with the cerebral cortex and cerebellum.

### Across modalities validation - The Richiardi list

[27] identified a list of 136 genes whose expression is correlated with the so-called resting-state functional connectivity [7]. We recomputed the Mapper network using the gene-list identified by [27] instead of the whole genome. We then compared this Mapper network to functional connectivity matrix derived from resting state fMRI data from the Human Connectome Project. In the Mapper generated networks we observe that again the cerebellum constitutes a stand-alone component, while the remaining samples are divided in two larger and anatomically coherent components: one containing most of the cortical samples and the other one containing mostly subcortical ones, see Fig. 5.

**Figure 5:**
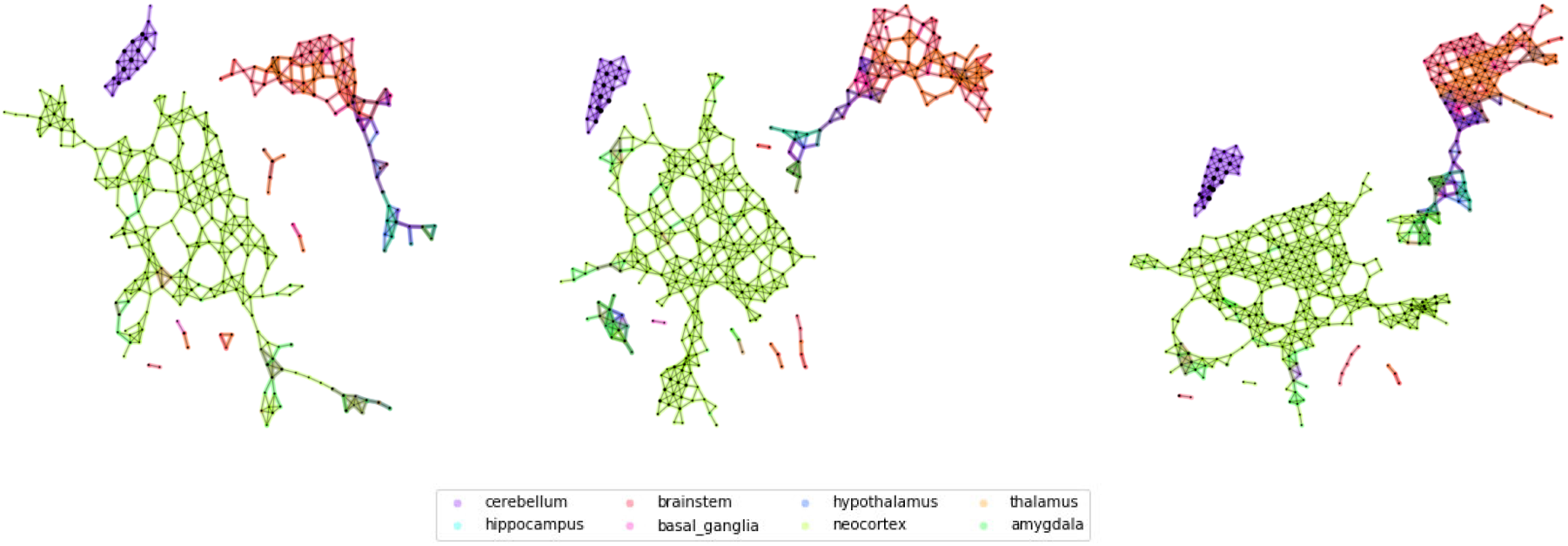
Mapper networks on reduced Richiardi functional list. From left to right, we show some Mapper networks obtained for different overlaps between bins (overlap 0.25,0.3, 0.35). The networks displays the separation of the cerebellum and two larger components, composed by cortical and subcortical areas repsectively, which we further characterize using the agreement matrices.

In this case we can compare the properties of the Mapper links by comparing them to the functional connectivity values. Indeed we find that the Mapper links correlate with higher functional connectivity. The links extracted by the mapper generated network are up to 14% more likely to include high functional connectivity between the ROIs than the complete full matrix, and up to 29% more likely than the ones ignored by the algorithm construction. These values were computed using the Kolmogorov-Smirnov statistic measuring the highest gap in the cumulative distributions, see Fig. 6e). Comparing more closely the distributions of functional connectivity for links presents and absent in the mapper network we can identify the level at which the change in trade-off occurs (Fig. 6d).

**Figure 6:**
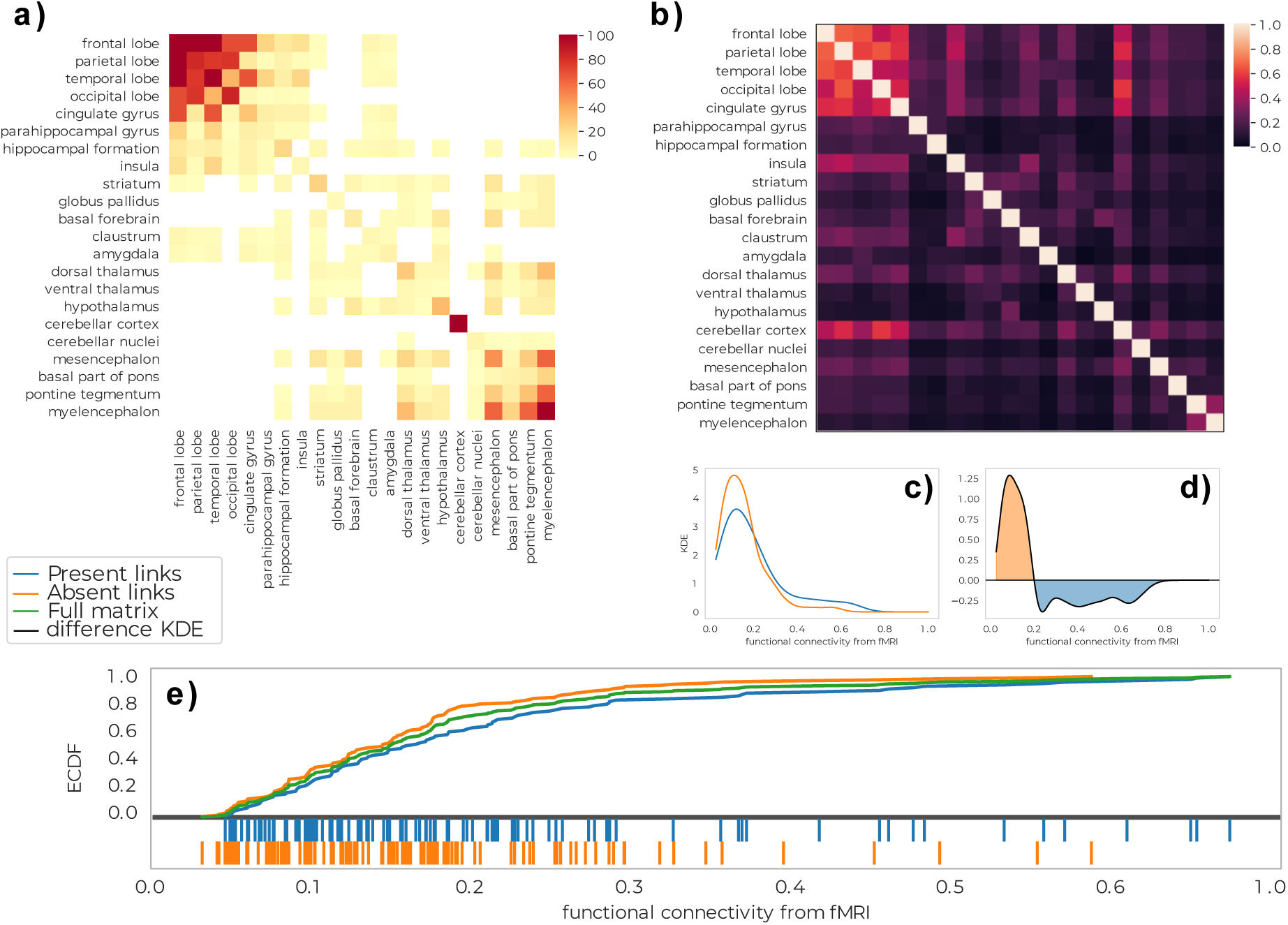
Comparison of the output network with an average functional network from fMRI. a) Agreement matrix between Mapper co-occurence matrices for Mapper networks built using Richiardi list. b) Average fMRI synchronization between regions. c) distributions of fMRI correlations for links that are and are not present in the Mapper networks. d) difference between the distributions in c). e) cumulative distributions.

In their work, [27] in order to remove highly differential outliers, decided to focus their analysis on 1777 cortex samples mappable via their Montreal Neurological Institute (MNI) coordinates to 13 functional networks (see S.I.for full list), excluding 1926 samples from the basal ganglia, cerebellum, and deep gray matter regions including the hippocampus.

As with the analysis of the AHBA whole-genome, we can use this information to define different regimes for the links and compare in more details the connectivity in the Mapper for the samples considered by [27] in their analysis and those ignored. The results are presented in Fig. 7.

**Figure 7:**
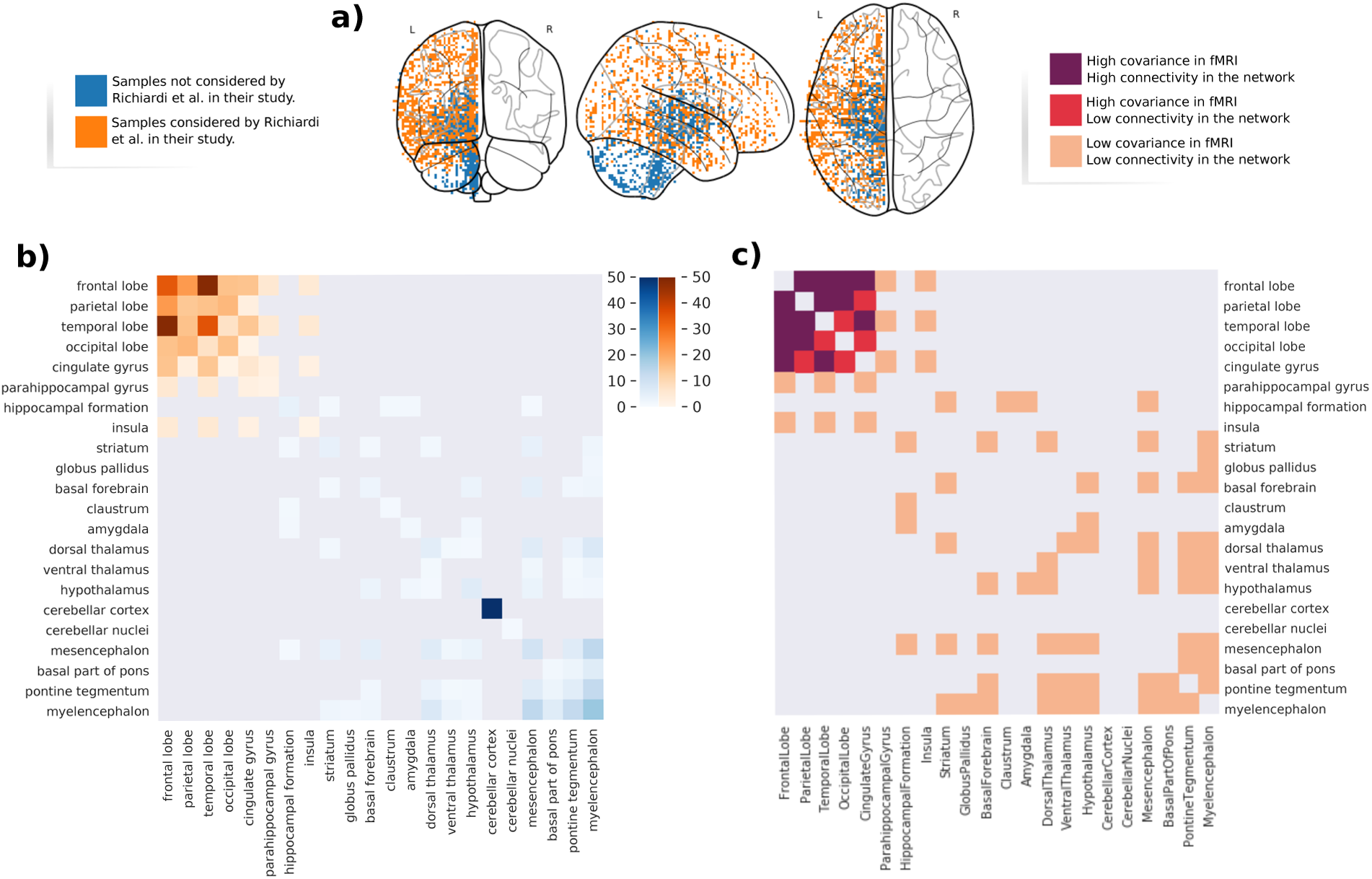
Comparison of the connectivity for the samples correlated with synchronous activity in fMRI and the rest of the brain. a) A visualization in the MNI space of the samples considered by [27] in their work (in orange) and the samples excluded in their work (in blue). b) The mapper agreement matrix reduced to contain only the co-occurances within the samples considered by [27] (in shades of orange) and the rest of the hemisphere (in shades of blue). c) the Agreement matrix colored according to three regimes based on the combinations of sparse/dense connectivity in the Mapper and low/high average functional connectivity.

When considering the connectivity of the ROIs in the network via the co-occurrence matrix we can see that there is a similar pattern to the one found in the fMRI covariance matrix where the ROIs involved in the functional networks studied by Richiardi. All the samples not considered by Richiardi in their work tend not to be clustered together in the Mapper network. Viceversa, the samples that Richiardi found to correlate with fMRI tend to be clustered together in the network, with the exception of the Temporal, Occipital lobes and Cingulate Gyrus which are less densly connected.

### Dopamine system

We can see two main connected components in the Mapper generated network, Fig.8 a). A small component containing only elements of the cerebellum and a giant component containing most of the samples. In the giant component two anatomically coherent modules are easily distinguishable: one containing samples from the cortex, the other one from the subcortical ROIs.

**Figure 8:**
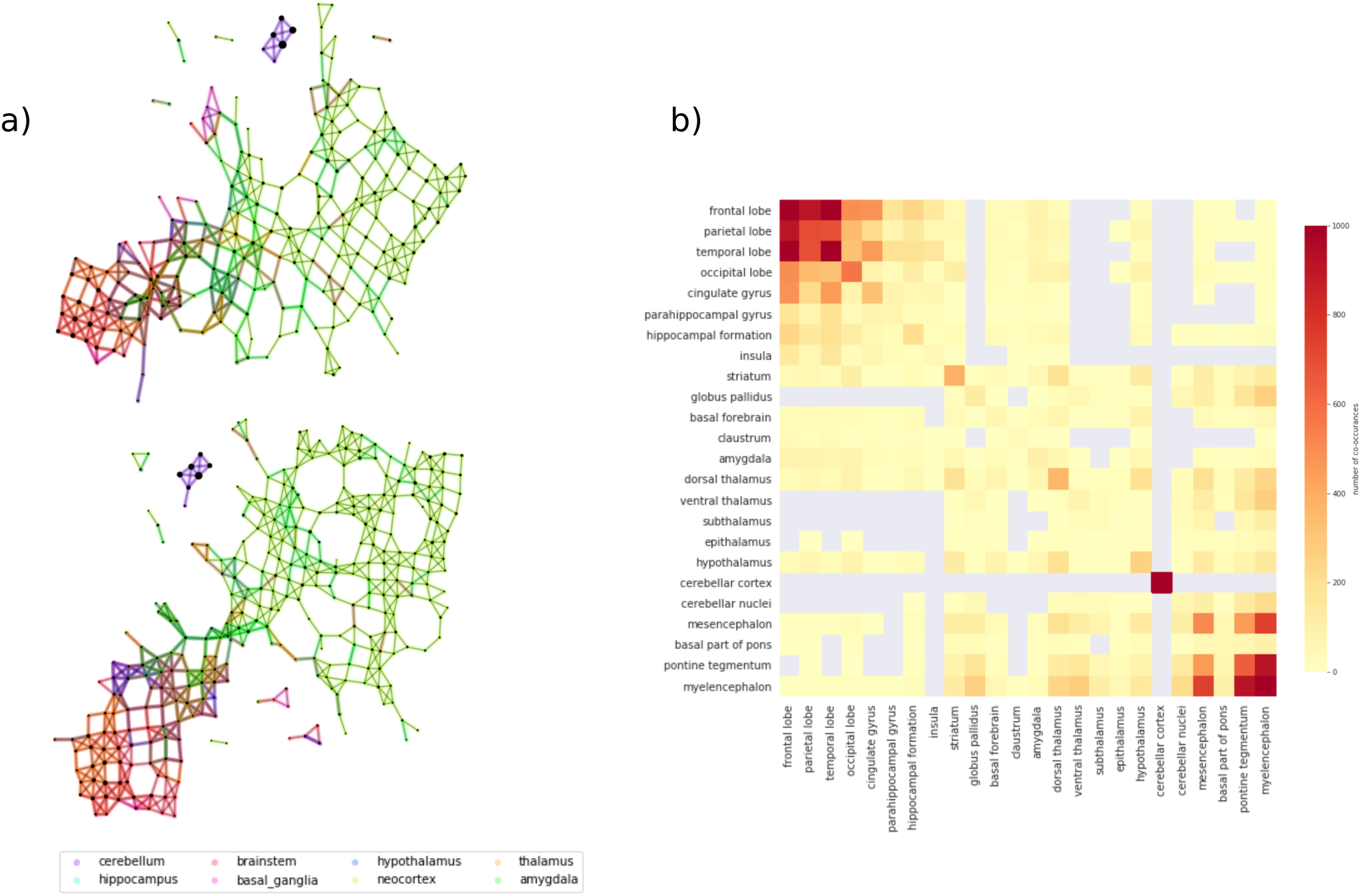
Mapper networks of the dopamine system. a) From top to bottom, we show some Mapper networks obtained for size 5%, with a 25% overlap (top), and size 6%, with a 20% overlap (bottom). The networks display very similar qualitative properties, e.g. the separation of the cerebellar areas, which we further characterize using the agreement matrix b).

From the agreement matrix we clearly distinguish the two modules as deep red blocks, Fig.8 b). The fact that a significant portion of the high-connectivity is between ROIs indicates that the clusters in the network are very inhomogeneous in their composition, i.e. it is very likely to find samples from different ROIs clustered together. The organization of the network in two modules is reminiscent of the anatomical organization of the mesolimbic dopamine pathway characterized by a crosstalk between cortical and sub-cortical structures. For this reason we decided to study the organization of the network relative to nodes containing samples from the substantia nigra and ventral tegmental area (VTA). These regions were chosen because they are the areas of the brain most densely populated by dopamine-producing neurons and therefore thought to be the starting point of the dopaminergic pathway in the brain [23] (see Fig. 9a). We then calculated the shortest path distance from these nodes to every other node in the main connected component. We discuss here the results for seeds chosen in the ventral tegmental area, but the analysis is consistent for the substantia nigra (see Fig. SI.5 for a comparison of the results between the two ROIs).

**Figure 9:**
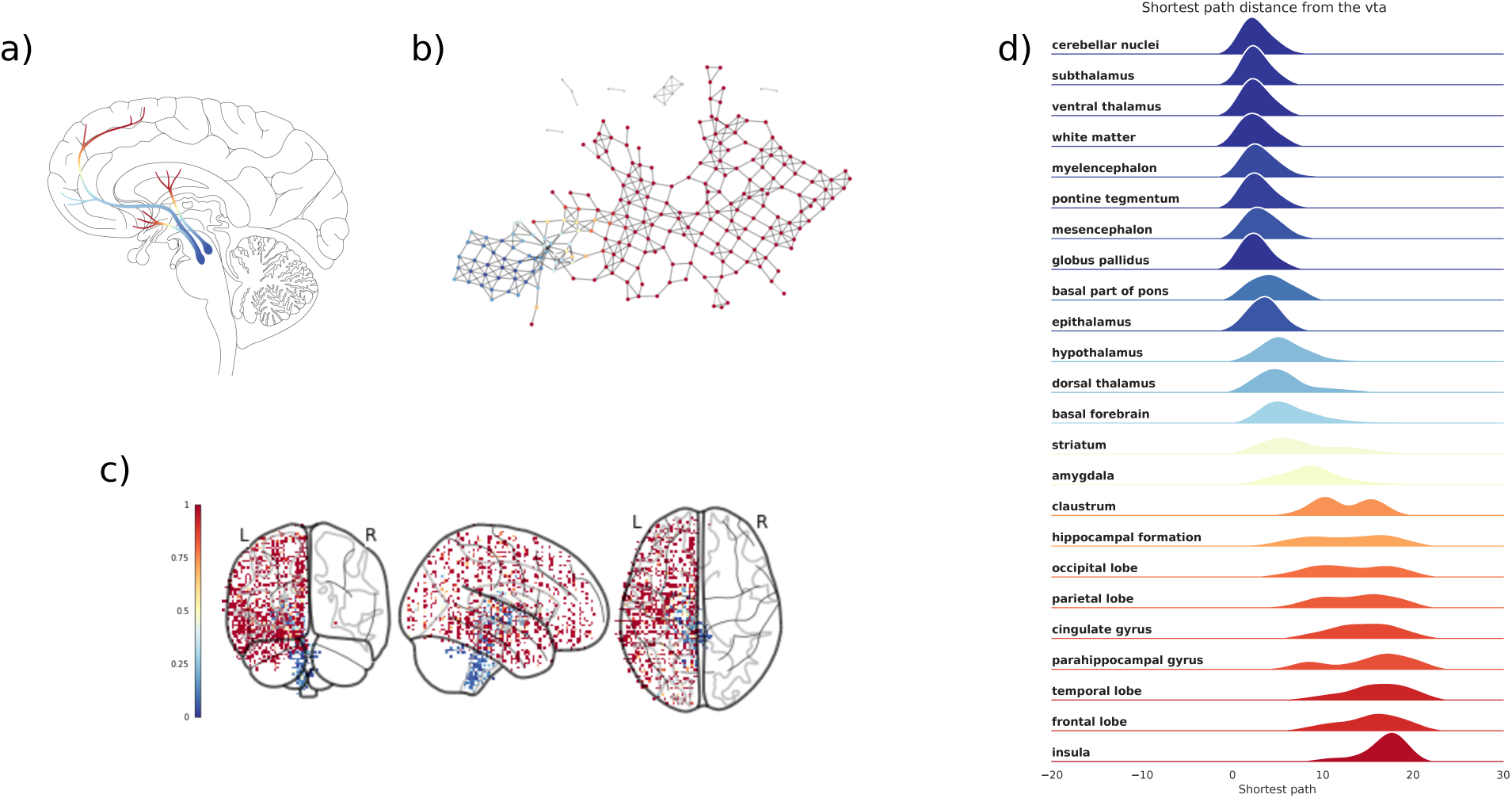
Distribution of shortest path distance in the output networks. For each network we identified the nodes in the network containing samples from the ventral tegmental area and computed the shortest path distance from these nodes to the rest of the network. a) The dopaminergic pathway (thanks to Gill Brown, London College of Communication, UK). b) Each node/cluster is colored according to its average path distance from nodes containing samples from the ventral tegmental area. c) The samples from the left hemisphere in the MNI space colored according to the average path distance of the nodes they belong to in the network. d) The distribution of path distance values for each ROIs ordered (top-bottom) according to their mean value from closest to farthest.

Each node/cluster thus had a value assigned to it that represents its distance in the Mapper network from the ventral tegmental area (Fig. 9b). Since each sample in the left hemisphere belongs to one or more nodes/clusters on in the network, the shortest path distance values was mapped from the nodes to the samples (Fig. 9c). We then studied the distribution of these network related values inside each ROIs (Fig. 9d). In blue the ROIs containing samples in the same cluster or closer to the substantia nigra. The samples from these ROIs are mostly contained in the same network community. The ROIs are ordered top to bottom from closest to farthest. In lighter blue and oranges the ROIs with samples in the part of the network bridging the two modules. In red the other module which encompasses most of the cortex. Interestingly, the ROIs in lighter blue and orange (e.g. the striatum and the thalamus) are actually ROIs that contain a high number of synapses between axons coming from the brainstem and neurons projecting into the cortex [25]. These results show that indeed the Mapper network reproduces closely the anatomical structure of the dopaminergic pathway.

## Discussion

Dimensionality reduction methods are as old as statistics and are an essential tool to study complex and complicated systems. However, what is gained in size is often lost in interpretability; it is therefore crucial to find lower dimensional representations of high dimensional spaces that are easily associated to other measurements or observables of the system under study. Among topological simplification tools [11, 26], Mapper is a method of choice to achieve this goal. Although a powerful tool that yielded significant results [20, 24, 31], little work has focused on the cross-validation of its results across modalities and data sets. In this study, we built a Mapper pipeline designed to extract pattern of similarity in genetic expression and crossed examined them through brain anatomy, with the aim to link genetic expression to neuroscience. Here we show that Mapper is an efficient topological simplification tool when applied to gene-expression data from the AHBA. Critically, it was able to extract from the data meaningful patterns of gene co-expression that are related to brain function and structure. First, we validated the pipeline by replicating most of the co-expression patterns obtained for the AHBA in [13, 14]. We found that in most cases the Mapper algorithm creates densely connected areas between samples that have a low differential gene expression such as areas within the neocortex. These results matched previous results [14] obtained with other network analytic techniques such as Weighted Gene Co-expression Analysis (WGCNA) [36]. However, the opposite does not hold for other areas such as the hippocampus and the amygdala. Clear outliers are samples from the amygdala which even with low differential expression tend to be overall sparsely connected. A possible explanation could be in the loadings of the first two principal components on these areas, motivation for the slicing into bins for local clustering. If the samples from the amygdala do not end up in neighboring bins it would be hard for them to cluster together. Indeed, a potential solution would be to adopt underlying filters based on recent dimensional reduction techniques designed to the effects of heterogeneous sampling of gene-space [21].

We are also able to reproduce the results obtained in [27], by correlating resting-state fMRI connectivity patterns with gene co-expression from a curated list of genes. This is extremely interesting as no specific choice was required to extract a topological manifold (the Mapper network) that linked directly gene expression to function. Moreover, recent results showed that the landscape of resting and task brain activations can be well approximated using Mapper on voxel-level activation data [31]. The homogeneity of the methods and descriptive spatial scales would then naturally allow to fuse the two approaches by characterizing the activity clusters at the genetic level and viceversa, or by producing brain activity Mappers informed by the underlying gene-expression Mapper networks.

Finally, armed with these validations, we turned to a subset of genes associated with a crucial neurostransmitter, dopamine. Remarkably, the Mapper network follows closely the anatomical dopaminergic pathway, elegantly relating complex genetic co-expression patterns to their physical manifestation. At this point, it is worth noting that the pre-processing of the data input to Mapper is minimal compared to traditional genetic studies [14, 17, 27] and that the results are stable across large parameter ranges, see S.I. and previous results [9, 10]. This is of great importance since the field is starting to raise concerns about reproducibility issues related with data pre-processing [2]. These results suggest that Mapper can be used to test hypothesis regarding the implication of specific genes or set of genes in brain function by producing associated networks and their relation with other imaging modalities such as EEG, MEG, fMRI, and potentially generic combinations of different modalities. This could provide an integrated representation of spatial, genetic and functional structure of the brain, and lead to direct applications in understanding the interactions and effect between neurotransmitters and finally shed light on the mechanism underlying i) mental health disorders and ii) the effect and side-effect of their associated treatments.

## Methods

To study the hidden organization of the gene-expression data set we use the Mapper algorithm [33]. The Mapper algorithm builds a low-dimensional skeleton of the data set using similarity information intrinsic to the original data set guided by other well established low-dimensional embedding techniques. In our work we set out to construct a skeleton that would represent the correlation similarity between the genetic expression of different brain regions.

### Mapper

The algorithm requires many parameters and choices that help build the network that best describes the aspect of the data set one wants to highlight. One can summarize the algorithm in 4 major parts (see Fig. 11 for a detailed description):

**Figure 11:**
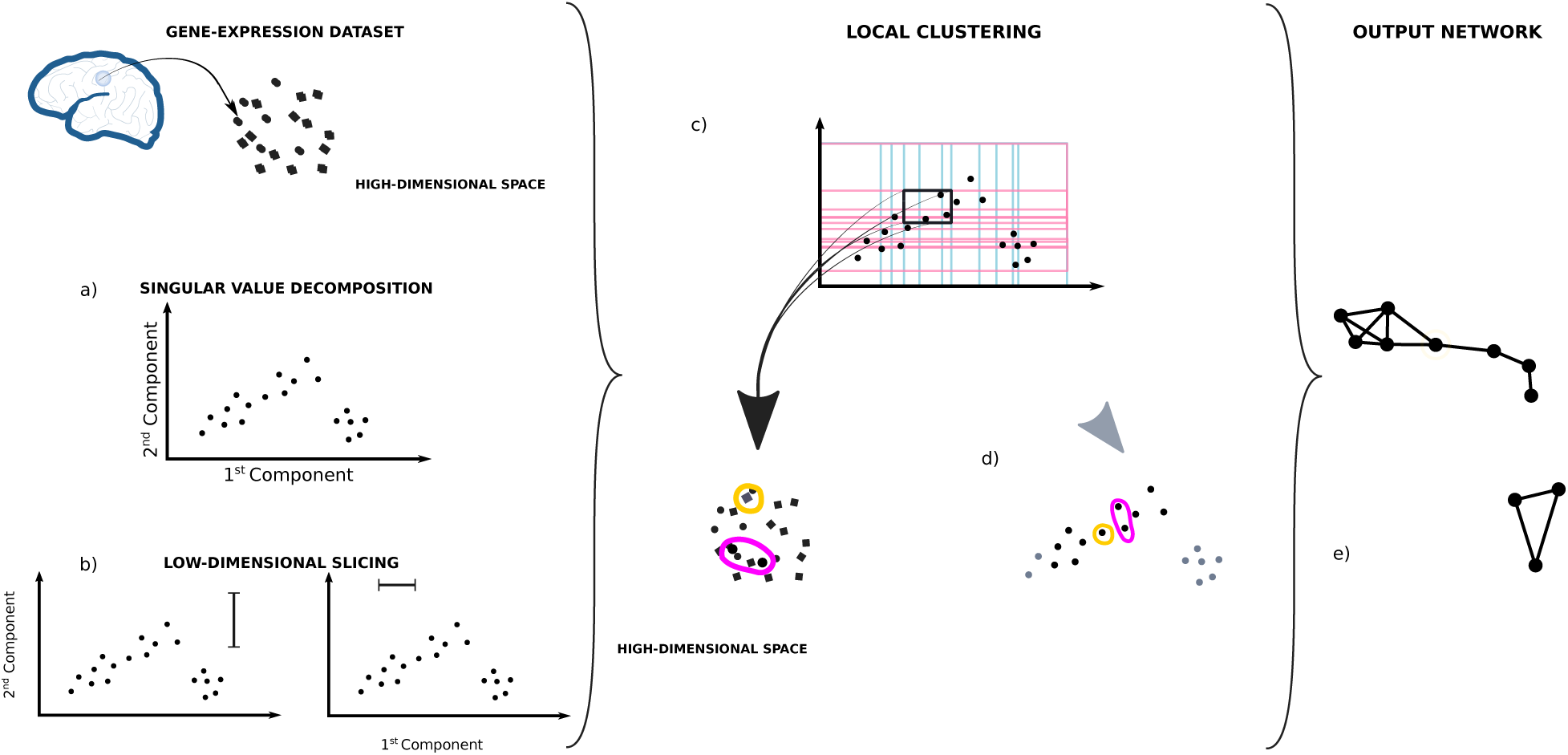
Description of the Mapper algorithm. Each point in the data-set is considered as a vector of gene-expression in a high-dimensional space (*∼* 29000 dim, 136 dim, or 56 dim according to the number of genes considered) a) The first two principal components are computed reducing the initial dimensions to 2, b) then the data set is sliced twice in overlapping windows of equal density along both components. c) The information from both slicing is used together to divide the data-set in overlapping bins. This way, each bin will have samples having similar weight in both component. d) The samples from each bin are clustered independently from each other according to the gene-expression correlation between the samples in the original high-dimensional space. e) The information is then summarized in a network were each independent cluster is represented by a node and two cluster are connected together if clustered belong to neighboring bins and share samples.

**Low-dimensional embedding** - A set of features of the data set are chosen as a local guide in the slicing process. In our case we chose the first 2 components of the singular value decomposition of the samples’ gene-expression covariance matrix.

**Low-dimensional slicing** - Each dimension of the embedding in 11a) is considered separately. The data set is divided using overlapping windows. The windows’ size and overlap are key parameters for the local topology of the output network.

**Local clustering** - Using the combined information of the slicing in 11b), the data set is divided in overlapping bins. A clustering algorithm is run in parallel on each bin independently. In our case we chose to run a density based clustering algorithm ([8]) with a correlation based similarity as distance.

**Building the network** - Each cluster found in 11c) is represented by a node in the network. Samples contained in the overlap between windows, will be present in more than one cluster. When this happens an edge is drawn between the representing nodes to depict the non-empty intersection.

The only two free parameters in our application of the algorithm are the window size and overlap at step b. Instead of choosing a single parameter we use basic network features (number of connected components and edge density) to get a set of optimal parameters and obtain a series of network descriptors at different level of coarseness (in S.I. we show the resulting optimal parameter for each list of genes).

#### Agreement Matrix

For each network, we can analyze the gene-expression similarity within anatomical brain regions via the study of node connectivity in the network. To summarize the anatomical information stored in each network connectivity we build a co-occurrences matrix, where each element *C_ij_* of the matrix represents the number of times that two samples from ROI *i* and ROI *j* are mapped in the same node/local cluster.

We condense the connectivity information from the networks built using all the optimal parameters in a unique matrix, where each element is the average co-occurance across all networks and *C_ij_* is non-zero only if ROI *i* and *j* are connected in all the networks.

It useful to notice that the co-occurance matrix can also be constructed considering only a subset of the samples present in the network. This approach gives a more selective account of the correlation between anatomical regions, where the influence of the ignored regions is still accounted for by the network but ignored in the numeric computation of the matrix. This effect can be noticed in Fig. 7 where the connectivity of the samples considered by [27] in their work where studied separately from the rest of the samples in the Allen human Brain Atlas.

## Supporting Information

### 5.1 Optimization of parameters for the Mapper algorithm

The Mapper method has 5 parameters that one needs to fix before running the algorithm:

**filter** the filter is a map, or series of maps, which will guide the subdivision of the data-set in overlapping bins.

We decided to use the first 2 principal components of the gene-expression covariance for the each list of genes.

**coarseness** the level of coarseness in the output network (number of nodes, threshold of connectivity) is determined by the size of the overlapping bins on which the clustering algorithm is performed. Big bins give fewer nodes than smaller ones. Bins with a high percentage of overlap will give a densely connected network with many nodes. Viceversa, lower overlap will give more connected components with few edges.

**window size**

**overlap**

**clustering** A clustering algorithm is run independently on each bin. The similarity measure used by this algorithm is the key to the interpretation of the output network.

**clustering algorithm** We chose HDBSCAN [8] to use a density based hierarchical algorithm that has a less supervised cutoff;

**similarity measure** Considering that we intended to study the relationship between gene-expressions, we chose 1*−*correlation between the gene-expression vectors.

#### 5.1.1 Find the window sizes that are going to give the most reasonable average bin size

We look at 50 different window sizes -from 1 to 50. The size is chosen so that a fix percentage of the data points are present in each window. To reduce the number of networks to build, we reduce the number of window sizes in consideration by studying the resulting average bin size. In this preliminary study the windows are built with no overlap. After intersecting the two sliced data-set, we obtain the list of all bins to consider for clustering. We compute the average bin size for each percentage value (see Fig.SI.1)

We chose window sizes that provide on average bin size of 5 data points or more. For window sizes that have the same average bin size, we chose the smallest and biggest percentage of points in the window. The resulting window sizes to study are:

[5, 6, 7, 8, 9, 10, 11, 12, 13, 14, 15, 16, 17, 19, 20, 21, 24, 25, 30, 33, 34, 40, 49]

We then construct the windows with these sizes with an overlap ranging between [5%, 85%] of the window size.

#### 5.1.2 Finding the best combination of window size to overlap

We study the 368 pairs of size and overlap using general structural parameters of the resulting networks. In particular we take into consideration connected components and edge density (ratio between existing edges and possible edges).

Here are the limitations that we considered:

STEP 1 - limiting the number of connected components: we impose to have more than 1 connected component that is not and isolated point - top plot and at least 50% of nodes in the graph to be isolated - bottom plot

STEP 2 - we limit the edge density of the connected components to 1. To do so, we assume the number of nodes in the lattice to be square and we can always consider that *N_c_*, the number of nodes in a connected component, is *n*^2^ *≤ N_c_ <* (*n* + 1)^2^. We chose to maintain the same inequality in the edge densities 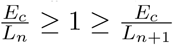 where *E_c_* is the number of edges in the connected component *c*, and *L_n_* is the number of edges in the square lattice with *n* nodes on each side.

After having imposed these two rules we are left with a few optimal parameters for our analysis.

**full** (window density, overlap) = [(5, 25), (5, 30),(5, 35), (6, 20)]

**fMRI correlated** (window density, overlap) =

[(5, 25), (5, 30), (5, 35), (5, 40), (6, 20), (6, 25), (7, 20)]

**dopamine** (window density, overlap) = [(5, 25), (6, 20)]

### 5.2 Defining the different regimes for comparison with existing results

For a more detailed comparison of the resulting mapper network connectivity with existing results found in the literature, we decided to divide the links found by the Mapper in 4 different regimes. The regimes were defined to highlight the relation between high/low connectivity in the mapper with high/low values in the comparing matrices (the differential expression matrix in Fig.3 and the functional connectivity matrix in Fig.6). In the figures SI.2 and SI.3 we show how the elements in the agreement matrix were divided in the 4 regimes respectively for the whole genome expression, and for the 136 genes identified in [27].

#### 5.2.1 Results from shortest path distance from ventral tegmental area

In Fig. SI.5 we show the distribution of shortest path distance from the substantia nigra and ventral tegmental area. The distributions represent the path distance from all samples in the seeded area to other ROIs present in the same component. ROIs with fewer samples or that are densely connected in the network have a narrower distribution. Overall the results for the two type of seeds are very similar to each other. This similarity is likely due to the samples from these two areas being clustered together in the same nodes in the network.

### 5.3 fMRI data acquisition and pre-processing

Subjects were scanned in a Siemens Connectome Skyra 3T scanner with a Gradient echo echo-planar imaging sequence (TR = 720 ms, TE = 33.1 ms, Kip angle = 52 degrees, multiband factor = 8, slice thickness = 2 mm, 72 slices and 1200 volumes) for the duration of 15 minutes. During the scan, the subjects fixed their gaze on a cross-hair on a black background. The images were subsequently processed in the following way. First, the structural pipeline was applied to create an undistorted native structural volume space for each subject. T1- and T2-weighted images were aligned, and bias field correction carried out. The pre-processed T1w were used for all subjects. Each subject’s native space was then registered to the MNI-152 2-mm space. The next step of segmentation was carried out in Freesurfer. The final part of structural pre-processing generated the NIFTI volume file, to which the functional data was mapped in the next step. This included functional pre- processing to remove spatial distortion, and realignment and registration to the structural MRI, bias field reduction, and normalization of the volumetric 4-D image to a global mean. Finally the 4-D data was masked with a final brain mask derived from a structural NIFTI file. No slice-timing correction was carried out, as the multiband factor required a large number of slices to be acquired close together. The ICA-FIX approach was then applied [32]. Motion correction was the next step of clean-up, using 24 confound time-series derived from motion estimation of the six rigid-body transformations, and their backwards-looking temporal derivatives. The resultant 12 regressors were squared. The motion parameters are subject to high-pass temporal filtering, and are then regressed out of the data.

### 5.4 List of 136 genes used for the fMRI comparison

[’ADAM23’, ‘ANKRD6’, ‘ATP6V1C2’, ‘BAIAP3’, ‘C3orf55’, ‘CARTPT’, ‘CCDC39’, ‘CD70’, ‘CDC2’, ‘CNTN6’, ‘CRYBA2’, ‘CTXN3’, ‘CXXC11’, ‘DMRT3’, ‘EPN3’, ‘FEZF1’, ‘FZD7’, ‘GAL’, ‘GLRA3’, ‘GNA14’, ‘GNGT2’, ‘GRP’, ‘HSD11B1’, ‘KANK4’, ‘KCNA1’, ‘KCNA3’, ‘KCNA5’, ‘KCNC1’, ‘KCTD15’, ‘KRT1’, ‘KRT31’, ‘LAIR2’, ‘LINC00238’, ‘LMOD3’, ‘LRRC38’, ‘LYPLA2’, ‘MGP’, ‘MYH7’, ‘MYLK3’, ‘NEB’, ‘NECAB2’, ‘NEFH’, ‘NEUROD6’, ‘NGFR’, ‘NOL4’, ‘NOV’, ‘NRP1’, ‘ONECUT2’, ‘PCP4’, ‘PIRT’, ‘PNMT’, ‘PRR15’, ‘PRSS23’, ‘PTGS1’, ‘RBP4’, ‘RBPMS2’, ‘RHOBTB2’, ‘RSPH9’, ‘SCARA5’, ‘SCN1B’, ‘SCN4B’, ‘SEMA7A’, ‘SHD’, ‘AC092324.1’, ‘SLC16A6’, ‘SLC22A10’, ‘SLN’, ‘SV2C’, ‘SYT10’, ‘SYT2’, ‘TDO2’, ‘TGFBI’, ‘PLAC2’, ‘TLX2’, ‘TNNT2’, ‘TRIM29’, ‘TSHZ3’, ‘AMDHD1’, ‘ASGR2’, ‘CD163L1’, ‘COL5A2’, ‘CYP2C18’, ‘FAM163A’, ‘GABRA5’, ‘GALP’, ‘GPR26’, ‘GPR88’, ‘GPX3’, ‘HPCAL1’, ‘IL13RA2’, ‘ISCU’, ‘NEXN’, ‘NKAIN4’, ‘NPBWR2’, ‘ONECUT3’, ‘OR51E2’, ‘PLCH1’, ‘PVALB’, ‘SNAP25’, ‘SPHKAP’, ‘TMEM52’, ‘TSPAN8’, ‘ZCCHC18’, ‘ALOX12’, ‘CALB1’, ‘CCBE1’, ‘CD6’, ‘CDR2L’, ‘CPLX1’, ‘ENPP6’, ‘GMPR’, ‘GOLT1A’, ‘GPR20’, ‘HOXD1’, ‘HPCA’, ‘IL33’, ‘IQCJ’, ‘KLK1’, ‘KLK8’, ‘LGR6’, ‘TCL1A’, ‘MS4A8B’, ‘MYBPC1’, ‘RP4-725G10.1’, ‘PYDC1’, ‘RTP1’, ‘SEMA3C’, ‘SH3RF2’, ‘SLC16A5’, ‘SLC20A2’, ‘SLC39A12’, ‘SOST’, ‘WISP1’, ‘WISP2’, ‘WNT4’]

### 5.5 List of 56 genes used for study on dopamine system

[’ADCY2’, ‘ADCY7’, ‘CDK5’, ‘CLIC6’, ‘COMT’, ‘DBH’, ‘DDC’, ‘DRD1’, ‘DRD2’, ‘DRD3’, ‘DRD4’, ‘DRD5’, ‘EPB41’, ‘EPB41L1’, ‘EPB41L2’, ‘EPB41L3’, ‘FLNA’, ‘GNAI1’, ‘GNAI2’, ‘GNAI3’, ‘GNAZ’, ‘GNB1’, ‘GNB2’, ‘GNB3’, ‘GNB4’, ‘GNG11’, ‘GNG3’, ‘GNG4’, ‘GNG8’, ‘KCNK3’, ‘KCNK9’, ‘MAOA’, ‘MAOB’, ‘NAAA’, ‘OC90’, ‘PPP1CA’, ‘PPP1CC’, ‘PPP1R1B’, ‘PRKACA’, ‘PRKACB’, ‘PRKACG’, ‘PRKAR2A’, ‘PRKAR2B’, ‘PRKX’, ‘PRKY’, ‘SLC18A2’, ‘SLC6A3’, ‘SNAP23’, ‘SNAP25’, ‘SNAP29’, ‘STX3’, ‘TH’, ‘VAMP1’, ‘VAMP2’, ‘VAMP3’, ‘VAMP8’]

### 5.6 List of functional Networks used in work by Richiardi et al

[’Auditory’, ‘dDMN’, ‘high Visual’, ‘Language’, ‘LECN’,’post Salience’, ‘Precuneus’, ‘prim Visual’, ‘RECN’, ‘Salience’,’Sensorimotor’, ‘vDMN’, ‘Visuospatial’, ‘Z restOfBrain’]

### 5.7 Code Availability

The code used for the analysis showed in this paper can be found at the following repository: https://github.com/alpatania/AHBA_microarray_Mapper/

## Acknowledgements

GP is supported by Compagnia San Paolo (ADnD Grant). This paper represents independent research part funded by the National Institute for Health Research (NIHR) Biomedical Research Centre at South London and Maudsley NHS Foundation Trust and King’s College London that support PS, OD, and MV. The views expressed are those of the authors and not necessarily those of the NHS, the NIHR or the Department of Health and Social Care. PS is supported by a PhD studentship jointly funded by the NIHR-BRC at SLaM and the Department of Neuroimaging, King’s College London. PE is supported by EPSRC through the EPSRC Centre for Mathematics of Precision Healthcare. fMRI data were provided by the WU-Minn HCP Consortium (principal investigators: David Van Essen and Kamil Ugurbil; Grant 1U54MH091657) funded by the 16 NIH Institutes and Centers that support the NIH Blueprint for Neuroscience Research and by the McDonnell Center for Systems Neuroscience at Washington University. Human microarray data are at http://human.brain-map.org. We would also like to acknowledge the contribution of Dr Gaia Rizzo (Invicro, London, UK) and Prof. Federico Turkheimer (Department of Neuroimaging, Institute of Psychiatry, Psychology and Neuroscience, King’s College London) for helpful discussion during the study set up. We would also like to thank Elene Kislitsyna (Department of Neuroimaging, Institute of Psychiatry, Psychology and Neuroscience, King’s College London) for her help on data analysis.

## Author contributions

Project was formulated by GP, PE, AP, PS and MV. PE and MV pre-processed the gene-expression data. OD processed functional connectivity data. AP implemented and ran the pipeline, and performed the analyses. AP, PS and MV interpreted the results from the study. All authors wrote the manuscript.

---

**Figure 10:**
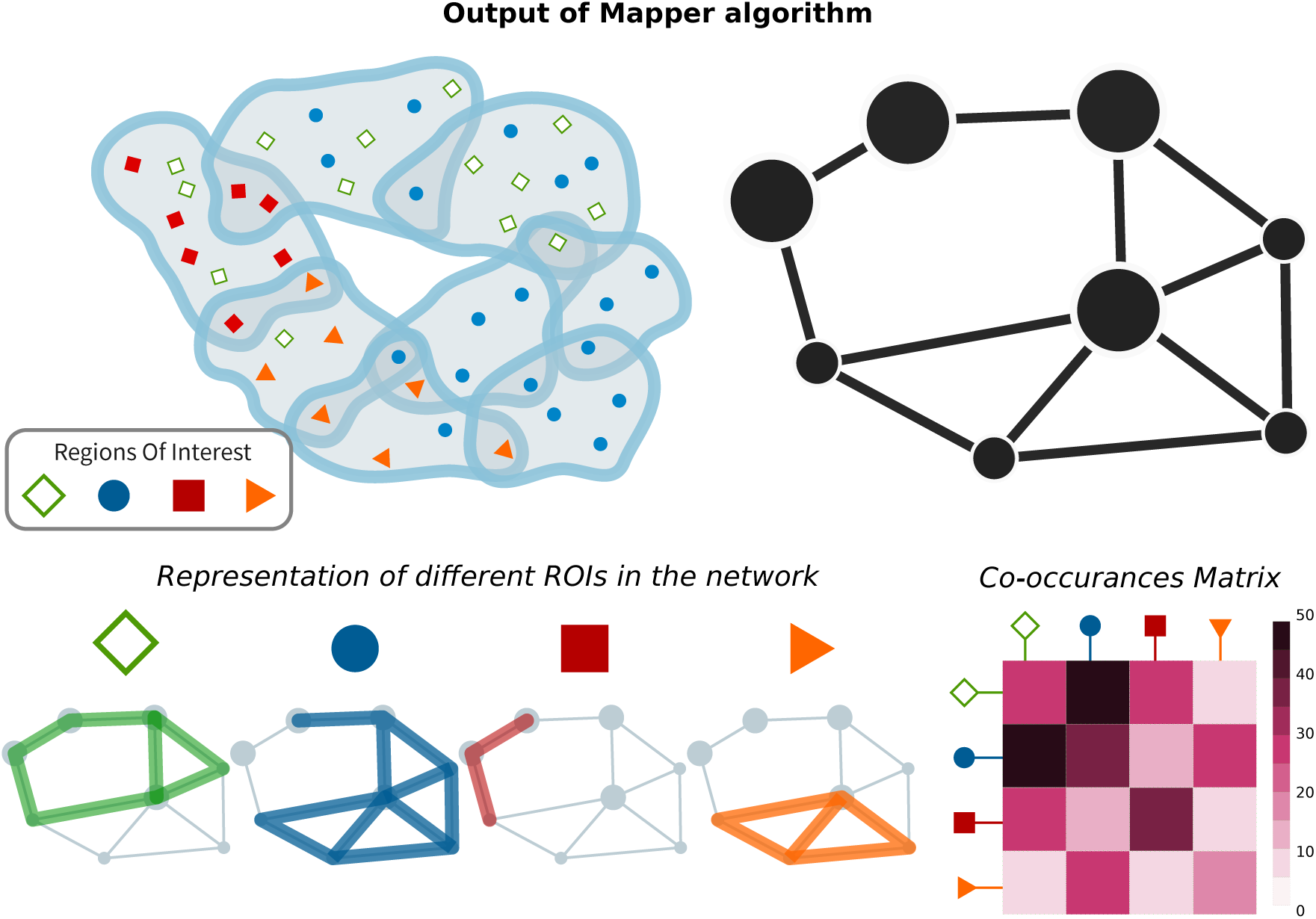
Comparison of the output network with differential analysis from the Allen Brain Institute. A) here goes the caption.

**Figure SI.1:**
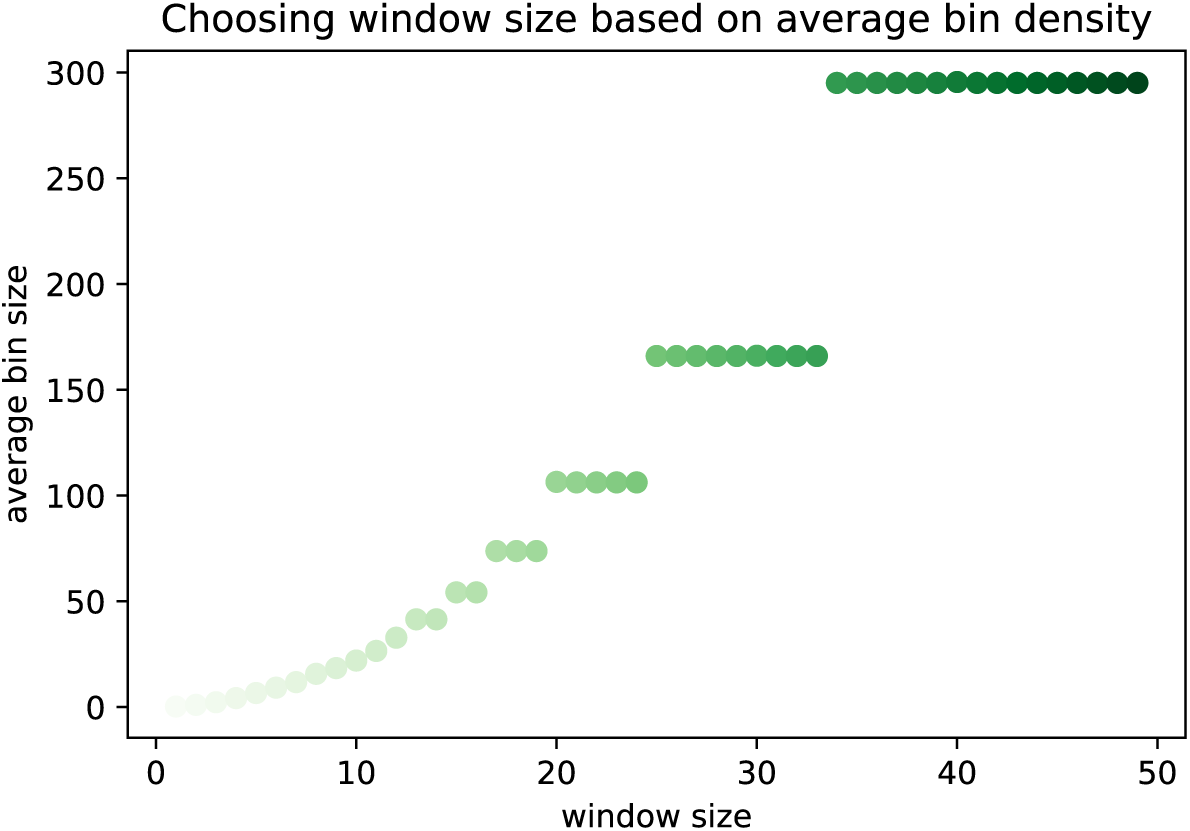
Average bin size as window density varies.

**Figure SI.2:**
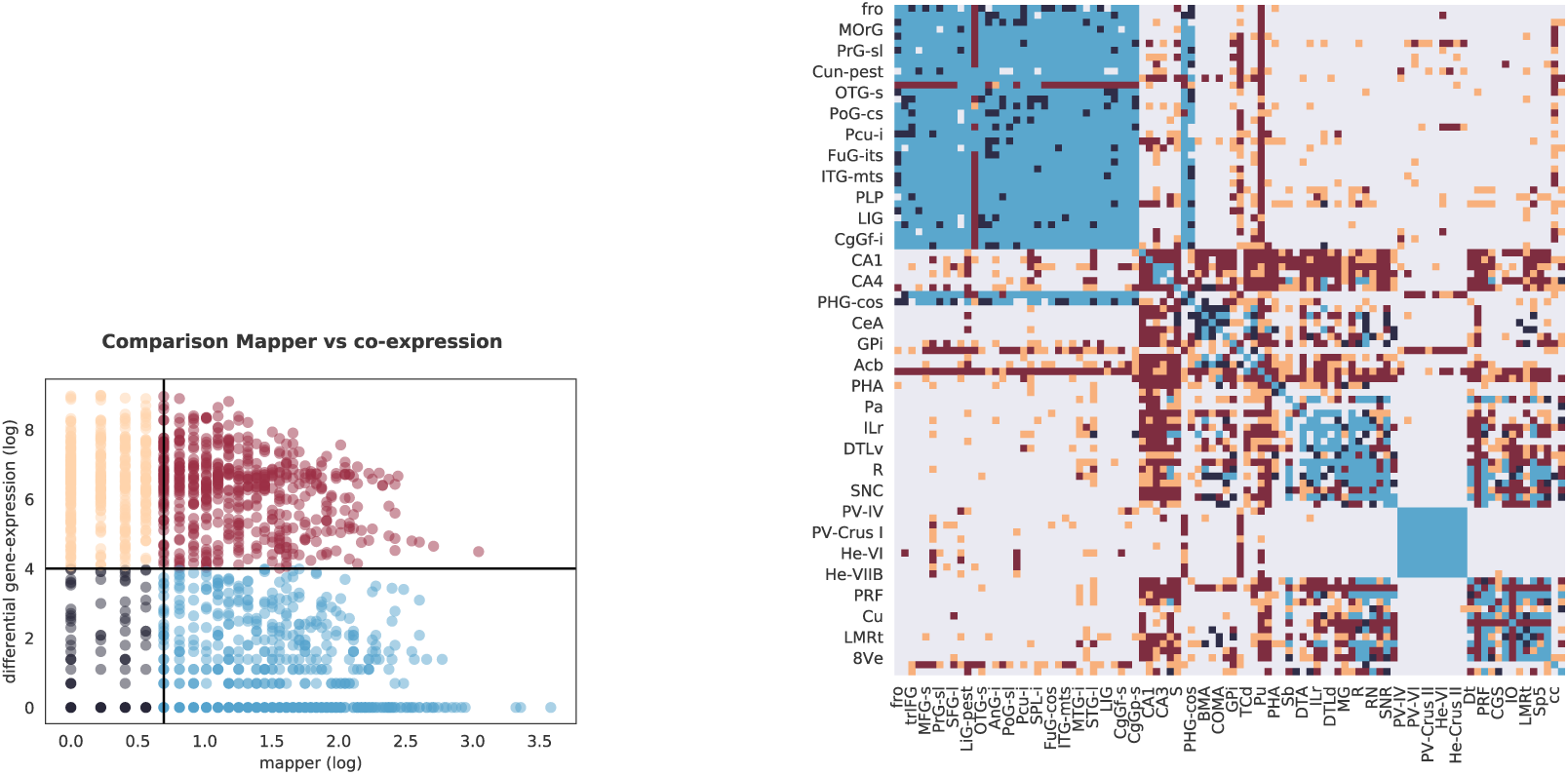
Defining connectivity regimes for whole transcriptome comparison. (left) Scatterplot depicting the subdivision of the links of the Mapper in the 4 different regimes. On the x axis the values of the agreement matrix divided between high/low connectivity at 2, on the y axis the differential expression of the links divided between high/low at 4, the value at which the difference between the distribution of differential expression in the present and excluded link changes sign 3d). (right) agreement matrix colored according to the 4 regimes to which each link belongs to.

**Figure SI.3:**
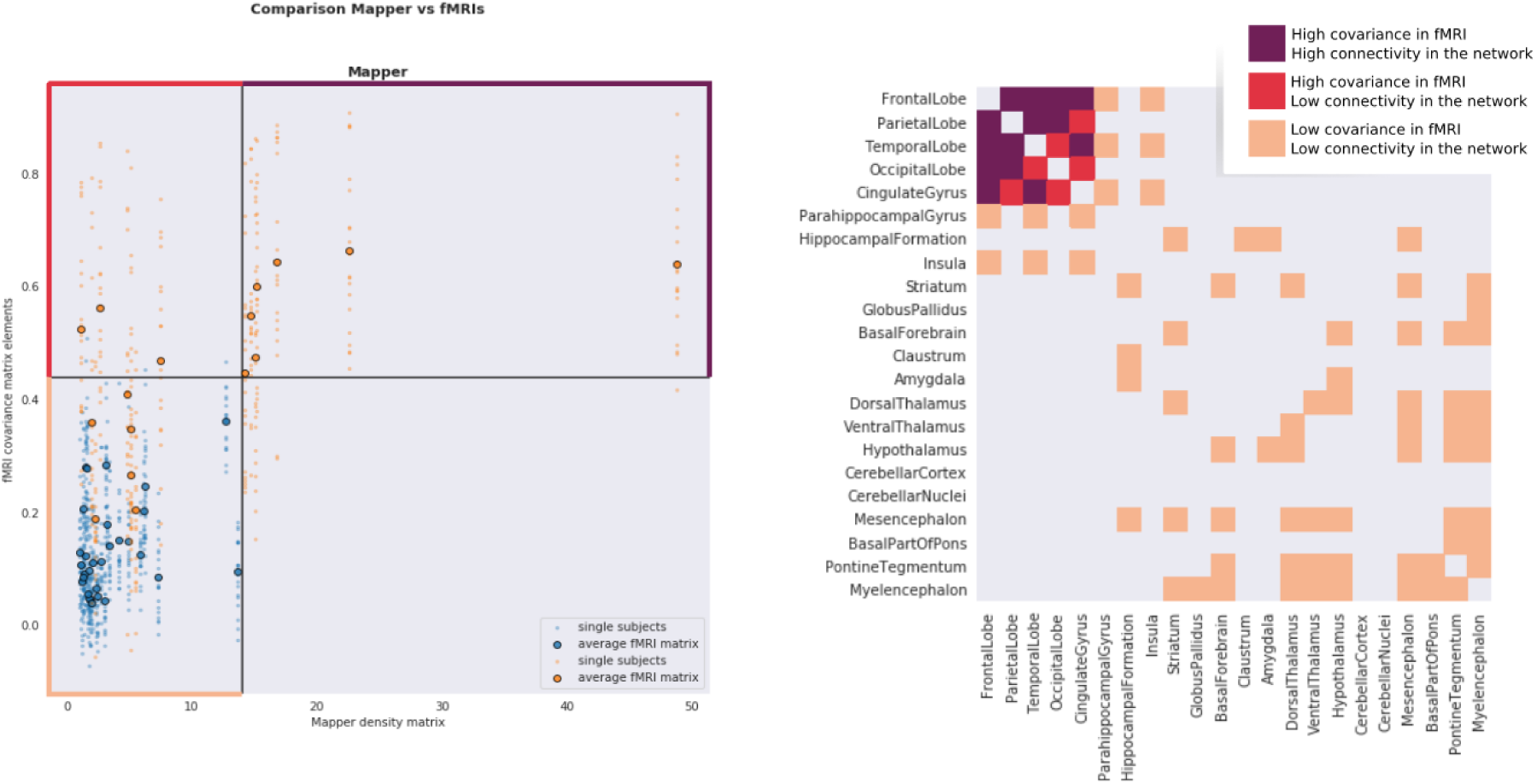
Defining connectivity regimes for fMRI comparison. (left) Scatterplot depicting the subdivision of the links of the Mapper in the 3 different regimes. On the x axis the values of the agreement matrix divided between high/low connectivity at 14, on the y axis the functional connectivity of the links divided between high/low at .44, the minimum functional connectivity value within the cerebral cortex (lobes and cyngulate gyrus). (right) agreement matrix colored according to the 4 regimes to which each link belongs to.

**Figure SI.4:**
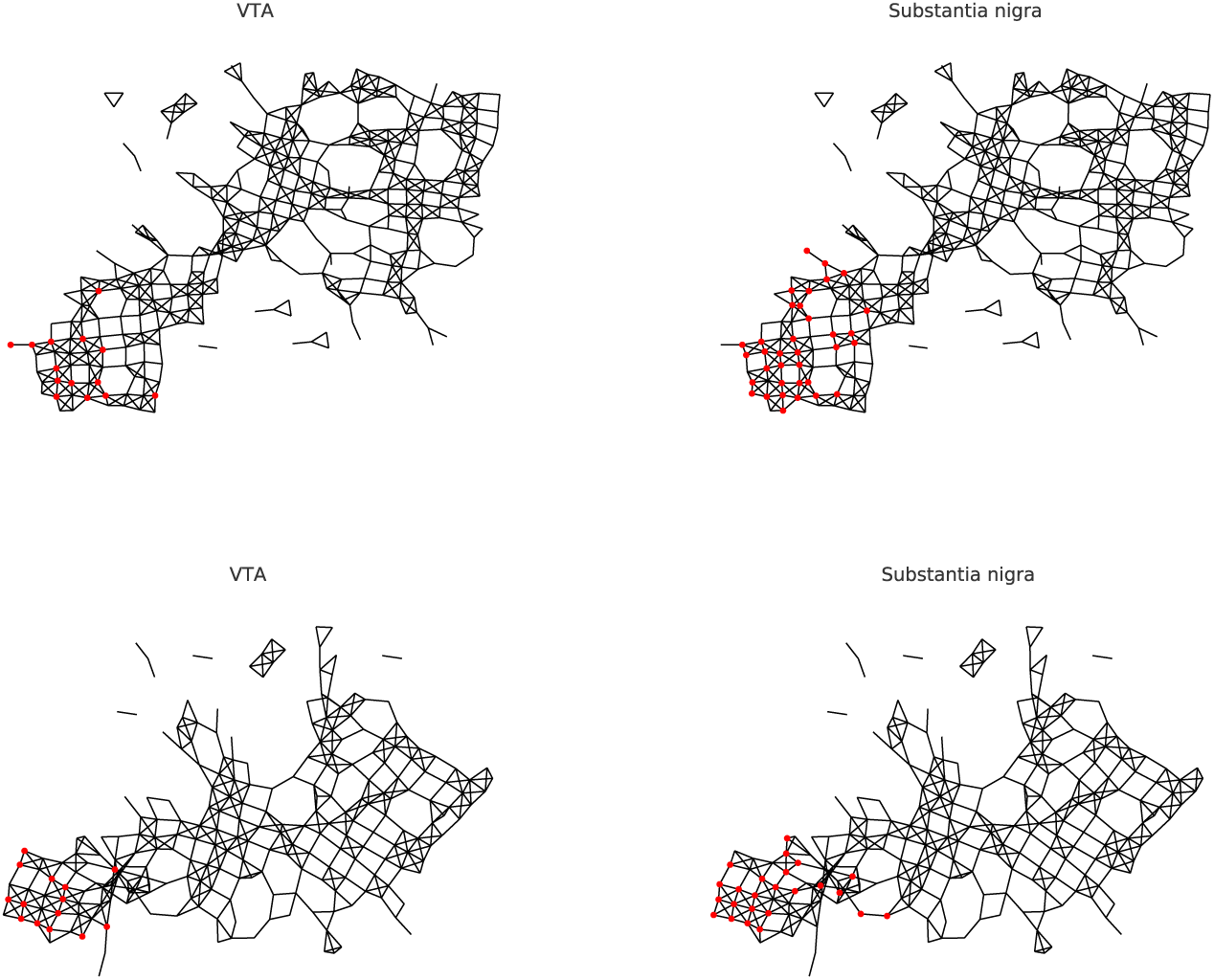
Placement in the networks of the seeds from VTA and Substantia nigra. For each network we identified the nodes in the network containing samples from the ventral tegmental area or substantia nigra and computed the shortest path distance from these nodes to the rest of the network. Here we show the nodes in network containg elements from the ventral tagmental area (left) and substantia nigra(right) for networks with parameters (size=5%, overlap= 25% (top) and (size=6%, overlap= 20% (bottom).

**Figure SI.5:**
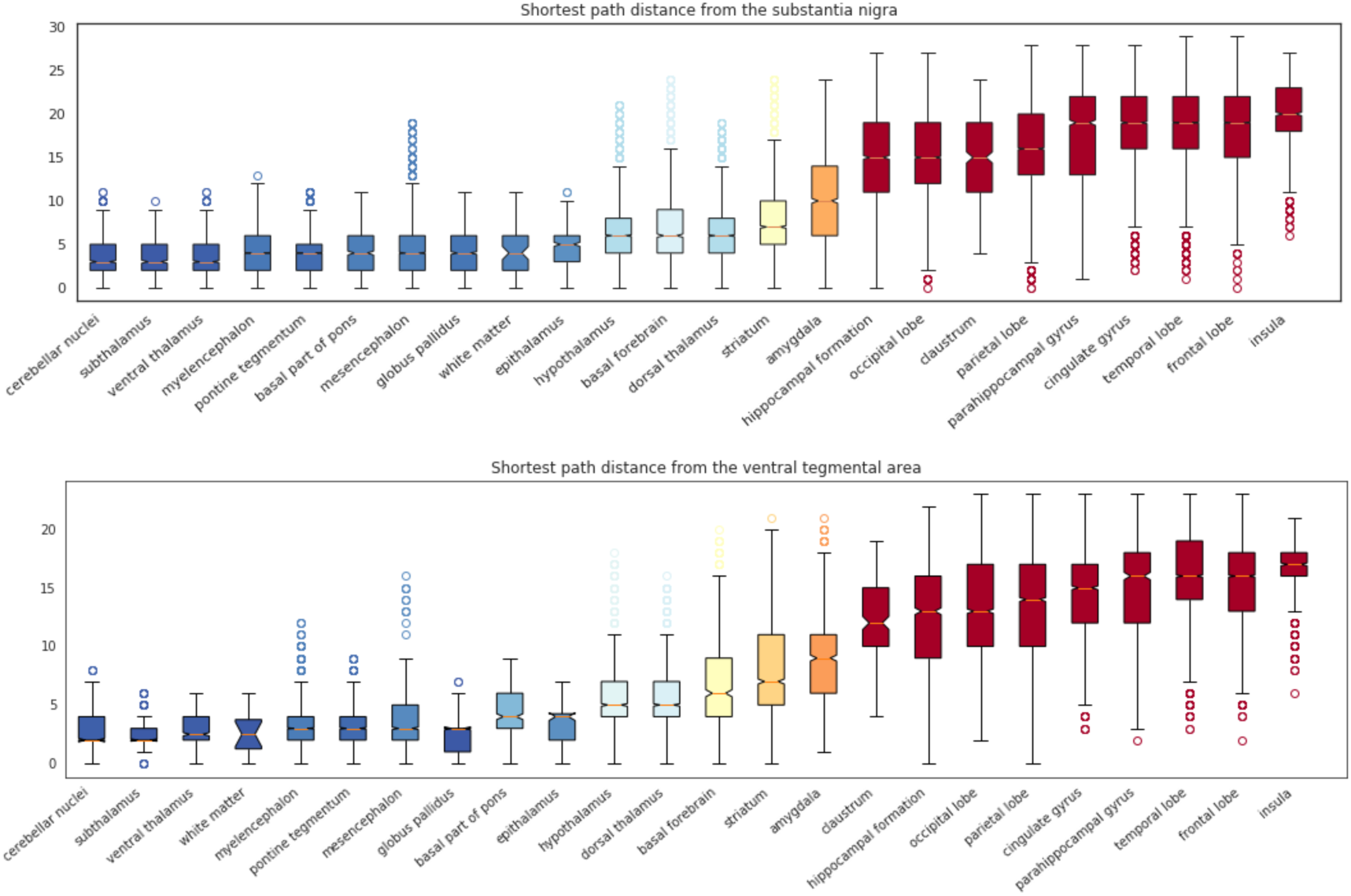
Comparison of the distributions of shortest path distance with seeds in VTA and Substantia nigra. For each network we identified the nodes in the network containing samples from the ventral tegmental area or substantia nigra and computed the shortest path distance from these nodes to the rest of the network. Here we show the distribution of path distance values for each ROIs ordered (left-right) according to their median value from closest to farthest, for seeds in the substantia nigra (top) and ventral tegmental area (bottom).

